# Ser/Thr Phosphorylation of *Mycobacterium tuberculosis* Type II Rel TA module by Protein Kinase K interferes with toxin neutralization: A novel mode of TA regulation

**DOI:** 10.1101/2023.12.15.571532

**Authors:** Shafinaz Rahman Sarah, Abhishek Garg, Shaleen Korch, Amita Gupta, Vandana Malhotra

**Affiliations:** Department of Biochemistry, Sri Venkateswara College, University of Delhi, New Delhi 110021, India; Department of Pharmacology, Midwestern University, Glendale, AZ, 85308, USA; Department of Biochemistry, University of Delhi South Campus, Dhaula Kuan, New Delhi 110021, India

**Keywords:** *Mycobacterium tuberculosis*, Toxin-Antitoxin modules, Persistence, Post-Translational Modification, Ser/Thr Phosphorylation, Regulation

## Abstract

Toxin-Antitoxin (TA) modules represent genetic elements implicated in bacterial persistence and antibiotic tolerance. Remarkably, *Mycobacterium tuberculosis* encodes 90+ TA modules, the majority of which are Type II TA comprising of a toxin component and an antitoxin counterpart that neutralizes the toxin. Upon exposure to stress, the antitoxin is degraded, releasing the toxin which then acts to halt cellular growth. Given that TA modules dictate social behavior of a population, we hypothesize that their regulation must be exquisitely controlled to avoid superfluous growth inhibition and initiation of persistence. However, the regulation and coordination of TA modules is poorly understood. Herein, we describe for the first time, a novel regulatory mechanism for Type II TA modules involving post-translational modification (PTM). Using computational tools, we observed that over 85 % of the *M. tuberculosis* TA proteins possess potential Ser/Thr phosphosites highlighting them as putative substrates for *M. tuberculosis* Ser/Thr protein kinases (STPK). We demonstrate that members of the RelBE family are subjected to *O-*phosphorylation by PknK, a stress-responsive growth regulatory STPK. Mass spectrometry confirmed multiple sites of PknK-mediated phosphorylation in the RelJK TA module. To gain insights into the functional impact of this PTM, we conducted *in vitro* binding and phenotypic growth studies with the wild type and mutant RelJK proteins. Our findings indicate that phosphorylation of Thr77 residue in RelK toxin compromises its binding to the RelJ antitoxin. These results suggest a potential role for *O-*phosphorylation in influencing the interaction dynamics of the TA module components.

**Importance:** Bacterial pathogens rely on the phenomenon of persistence as a survival strategy to combat the adverse environmental conditions encountered during infection. As a stochastic process, the driving force(s) that potentiate the formation of persisters in a bacterial population are largely unclear. This study is a step towards the discovery of intricate regulatory mechanisms that coordinate a synchronized TA cellular program. We propose a model wherein the TA module is regulated post translationally, specifically via Ser/Thr phosphorylation disrupting the interaction between the toxin and antitoxin proteins as a mechanism to regulate TA function.

## Introduction

Tuberculosis (TB) originated as a devastating disease in western Europe during the industrial revolution and remains a serious infectious disease accounting for 1.6 million deaths worldwide (1). The continued pathogenic success of *Mycobacterium tuberculosis,* the causative agent of TB, can be attributed to its ability to adapt and persist in host granulomas, in a non-replicating and drug tolerant state for decades with the potential of disease reactivation in elderly or immunocompromised individuals (2, 3). While considerable progress has been achieved in TB research over the last two decades, enhancing our comprehension of host-pathogen interactions, immune responses, and shifts in metabolic processes during infection, the precise mechanisms that drive the adaptive strategies of mycobacterial growth remain elusive.

By definition, a persister cell is one that displays a remarkable ability to tolerate adverse conditions and survive exposure to antibiotics or other stress-inducing factors that would typically kill the majority of the bacterial population (4–6). Physiological heterogeneity of a bacterial population enables the formation of persisters in a probabilistic manner. Numerous studies predominantly using *E. coli* as model bacteria, have revealed a variety of factors and stress response pathways that seem to be connected to the development of persister cells (7–11). One such means of achieving persistence involves toxin-antitoxin (TA) modules that consists of a pair of closely linked genes encoding a toxin and an antitoxin (12, 13). The toxin component of the module is typically a protein that, when present in excess, interferes with essential cellular processes, resulting in growth arrest or cell death (13). Conversely, the antitoxin is a protein or RNA molecule that counteracts the toxic effects of the toxin by binding to it, sequestering it, or inhibiting its biological activity (14, 15). Based on the nature of the antitoxin, eight families of TA modules have been described (16) with Type II TA modules representing the most widely studied and abundant TA family observed in prokaryotes. Typically, Type II modules are composed of a protein toxin and a cognate antitoxin protein that interact via protein-protein interaction (17). While several Type II systems have been evaluated in *E. coli* and *M. smegmatis* for their cytotoxicity (18–21) others such as VapBC, MazEF, ParDE, RelBE and HigAB have been implicated in stress-induced survival of *M. tuberculosis* (19, 22–25).

Understanding how TA expression is controlled and fine-tuned is critical to our comprehension of their role in mycobacterial physiology, fitness and pathogenicity. In general, most TAs are autoregulated by either the antitoxin protein alone, or by the toxin-antitoxin complex (12). Understandably, the relative stoichiometry of the toxin and antitoxin proteins are crucial determinants of their interaction outcomes. It follows that the coordinated expression of TA modules and regulation of toxin:antitoxin binding, and antitoxin degradation must be precise, and likely dependent on multiple factors such as the binding affinities of the toxin and antitoxin, stability of small antisense RNA, and post-translational modifications (26–28). A recent report describing toxin neutralization by its cognate antitoxin (or atypical kinase) via phosphorylation has generated significant interest (29). Likewise, Dawson et al., demonstrated post-transcriptional regulation of the *M. tuberculosis* RelBE system (30). Given their crucial roles in cell death and persistence, we are still far from decoding how TA modules are regulated, both as independent units, and as multi-family complexes.

Post-translational modification (PTM) such as *O-*Phosphorylation, also known as Ser/Thr/Tyr phosphorylation is one of the most universal forms of regulating protein function by covalent modification (31). These PTMs allow the bacteria to rapidly respond to changing environments and adapt to various conditions making them crucial for bacterial physiology and survival. The *M. tuberculosis* genome encodes eleven “eukaryotic-like” Ser/Thr protein kinases (STPK) and three phosphatases that help regulate the extent and duration of *O*-phosphorylation (32, 33). The reversible nature of phospho-Ser/Thr/Tyr signaling along with the wide array of diverse functions governed by *O*-phosphorylation, strengthens its role as a prominent central regulatory mechanism.

In this study, we investigated the regulation of TA modules by post-translational modification such as Ser/Thr phosphorylation. Given our previous studies on TA modules, we focused on the *M. tuberculosis* RelBE TA family that includes three TA loci namely, RelBE (*Rv1246c-1247c)*, RelBE2 (*Rv2865-Rv2866)* and RelBE3 (*Rv3357-Rv3358).* Since these were originally referred to as RelBE, RelFG and RelJK, respectively (19), we maintained the same nomenclature in this study. All three loci have been characterized as bona fide TA modules (19) that are responsive to stress environments such as nitrogen starvation and inhibit growth by targeting translation (34). We explored a link between the Rel TA modules and PknK, a cytosolic Ser/Thr Protein Kinase (STPK) that is coincidentally also induced under nitrogen starvation (35), and is implicated in translational control mechanisms (36). We demonstrate that PknK interacts with the RelE toxin and RelJK TA proteins *in vivo*. LC-MS/MS analysis identified several sites in the RelJ antitoxin and one site (Thr77) in the RelK toxin as sites of PknK-mediated phosphorylation. Co-expression and *in vitro* binding studies of the RelJK_WT_ and RelJK_Mut_T77A proteins in *E. coli* highlight a role for Thr77 residue in enabling stable toxin-antitoxin interaction such that its phosphorylation disrupts its interaction with the RelJ antitoxin. These findings reveal a unique regulatory mechanism to modulate TA function.

## Materials and Methods

### Bacterial strains, plasmids and culture conditions

For cloning and expression of recombinant proteins, *E. coli* DH5ɑ and *E. coli* BL21 (DE3) strains have been used respectively. *E. coli* was routinely grown in LB broth or LB agar plates at 37°C or in 2xYT medium for large scale protein purification. For Mycobacterium-Protein Fragment Complementation assay (M-PFC), *Mycobacterium smegmatis* transformants were plated on Middlebrook 7H11 agar (Difco) supplemented with 0.5 % glycerol, 0.5 % glucose and 0.2 % Tween and grown at 37°C. The cultures were supplemented with antibiotics or inducing supplements at the following concentrations: Ampicillin (Amp), 100 µg/mL; Kanamycin (Kan), 50 µg/mL; Hygromycin (Hyg), 50 µg/mL; Trimethoprim (TRIM), 25 µg/mL; IPTG (isopropyl-β-D-thiogalactopyranoside), 1 mM; Glucose (Glc), 1 %; Arabinose (Ara), 0.2 %; and Anhydrotetracycline (ATc), 50 ng/mL. All chemicals were procured from Sigma-Aldrich, USA unless stated otherwise. Details about strains and plasmids can be obtained upon request from V. Malhotra.

### Mycobacterium protein fragment complementation assay

To assess whether Rel toxins or antitoxins interact with PknK *in vivo*, the mycobacterial protein fragment complementation (M-PFC) system was used (37). Briefly, when two mycobacterial interacting proteins are independently fused with domains of murine dihydrofolate reductase (mDHFR), functional reconstitution of the two mDHFR domains can occur in mycobacteria, allowing for the selection of mycobacterial resistance against TRIM. *M. tuberculosis relB, relF or relJ* genes cloned into pUAB100 and pUAB300, episomal mycobacterium-*E. coli* shuttle plasmids gave rise to fusions of the [F1,2] mDHFR domains to each antitoxin (i.e., RelB_[F1,2]_). *M. tuberculosis relE, relG or relK* genes cloned into the integrating mycobacterium-*E. coli* shuttle plasmids pUAB200 and pUAB400 resulted in the fusion of the [F3] mDHFR domain to each toxin (i.e., RelE_[F3]_). Plasmids pUAB100 and pUAB200 generate C-terminal fusions (subscript “-C”), whereas pUAB300 and pUAB400 plasmids generate N-terminal fusions (subscript “-N”). Full length *pknK* cloned into pUAB300 and pUAB400, resulted in PknK_[F1,2]-N_ and PknK_[F3]-N_.

For all pUAB100 and pUAB200 clones, the GCN4 domains from pUAB100 and pUAB200 were replaced with *rel* DNA sequences. *Mycobacterium smegmatis (M. smegmatis)* mc^2^155 was co-transformed with plasmid constructs that encoded a toxin or antitoxin and *pknK*. As negative controls, a pUAB plasmid containing *relB* or *relE* was co-transformed with an empty vector (i.e., RelB_[F1,2]-C_/pUAB200), whereas a positive control was provided by the GCN4_[F1,2]_/GCN4_[F3]_ (GCN4 leucine zipper, *Saccharomyces cerevisiae* interacting domains in pUAB100 and pUAB200) (38). All transformants were plated on 7H11-Kan-Hyg plates and incubated at 37°C for 3 days. Transformants were restreaked onto 7H11-Kan-Hyg and 7H11-Kan-Hyg-TRIM and then incubated at 37°C for 3 to 5 days.

### Cloning, expression and purification of Rel TA and PknK proteins

*M. tuberculosis* genomic DNA was used as a template to amplify the *rel* genes: *relBE*, *relFG*, and *relJK*. The amplified products were digested with NdeI and Bsu36I restriction enzymes and cloned in IPTG-inducible expression vector pVLExp4231 digested with the same restriction enzymes. The clones were checked by restriction digestion and DNA sequencing. The wild type and mutant PknK and Rel TA proteins were purified as recombinant His-tagged proteins by Ni-NTA affinity chromatography. While RelB, RelF and RelJ antitoxin proteins were purified under native conditions, the wild type toxins (RelE, RelG, RelK) and RelK_Mut_T77A protein formed inclusion bodies and thus, were purified under denaturing conditions followed by on-column refolding using standard protocols.

### *In vitro* kinase assay

Purified wild type PknK protein kinase (1 µM) and substrate proteins (30 µM) were mixed in a tube containing 1x kinase buffer (25 mM HEPES pH 7.4, 15 mM MgCl_2_, 5 mM MnCl_2_, 1 mM DTT, 1 mM ATP) and incubated at 30°C for 30 minutes. The reaction was stopped using 5x SDS sample loading dye, followed by SDS PAGE for separation of proteins. After electrophoresis, the gel was processed for staining by ProQ™ Diamond Phosphoprotein Gel Stain (Thermo Fisher Scientific™) as per manufacturer’s protocol. The ProQ-stained gels were scanned with Typhoon Trio Variable Mode imager (GE Healthcare). The gel was then stained with Coomassie Brilliant Blue R250 (CBB R250) to visualize the protein bands.

Kinetic assays were performed to follow PknK-mediated phosphorylation of Rel toxin and antitoxin proteins over time. Purified RelE, RelJ and RelK proteins were incubated with active PknK at 30°C for 0, 15, 30, 45 and 60 minutes. Kinase assay was performed as described above and relative phosphorylation activity as a function of time was calculated using ImageJ software. To assess signal stability, kinase assay reactions were put up to allow maximal phosphorylation followed by desalting to remove ATP and allowed further incubation till 24 hrs. The reactions were subsequently analyzed by SDS PAGE.

### Mass spectrometry

For identification of phosphorylated residues, *in vitro* kinase assays were performed as described above and the proteins were resolved by SDS-PAGE. Individual proteins alone were processed similarly as negative controls. The gel bands observed after CBB staining were excised and subjected to LC-MS/MS analysis. Mass spectrometry was performed by Valerian Chem Pvt. Ltd., New Delhi as follows; **Sample preparation**: Gel bands were cut into small pieces and reduced with 5 mM TCEP and further alkylated with 50 mM iodoacetamide and then digested with Trypsin (1:50, Trypsin/lysate ratio) for 16 h at 37°C. Digests were cleaned using a C18 silica cartridge to remove the salt and dried using a speed vac. The dried pellet was resuspended in buffer A (2 % acetonitrile, 0.1 % formic acid), **Mass spectrometric analysis of peptide mixtures:** Experiments were performed on an Easy-nlc-1000 system coupled with an Orbitrap Exploris mass spectrometer. 1µg of peptide sample were loaded on C18 column 15 cm, 3.0μm Acclaim PepMap (Thermo Fisher Scientific™) and separated with a 0–40 % gradient of buffer B (80 % acetonitrile, 0.1 % formic acid) at a flow rate of 300 nL/min) and injected for MS analysis. LC gradients were run for 110 minutes. MS1 spectra were acquired in the Orbitrap (Max IT = 60 ms, AGQ target = 300 %; RF Lens = 70 %; R=60K, mass range = 375−1500; Profile data). Dynamic exclusion was employed for 30s excluding all charge states for a given precursor. MS2 spectra were collected for top 20 peptides. MS2 (Max IT= 60 ms, R= 15K, AGC target 100 %), **Data processing:** All samples were processed, and RAW files generated were analyzed with Proteome Discoverer (v2.5) against the Uniprot *M. tuberculosis* H37Rv database. For dual Sequest and Amanda search, the precursor and fragment mass tolerances were set at 10 ppm and 0.02 Da, respectively. The protease used to generate peptides, i.e., enzyme specificity was set for trypsin/P (cleavage at the C terminus of “K/R: unless followed by “P”). Carbamidomethyl on cysteine as fixed modification and oxidation of methionine and N-terminal acetylation were considered as variable modifications for database search. Both peptide spectrum match and protein false discovery rate were set to 0.01 FDR.

### Site-directed mutagenesis

For mutagenesis of RelK, threonine at position 77 was replaced with alanine using the primer pairs as follows: Forward Primer; 5’-CGACGAAGTCGCGATGCTGAAG-3’ and Reverse Primer; 5’-TCGCCCGCTCGATACACC-3’. Site-directed mutagenesis (SDM) was performed with Phusion™ SDM Kit (Thermo Fisher Scientific™) to generate the expression vectors, pVLExp::*relK*T77A and pCAK::*relK*T77A. The clones containing the desired mutation were confirmed by DNA sequencing.

### Phenotypic growth analysis

*E. coli* BL21 (DE3) cells were transformed with the expression vector pVLExp::*relK* and pVLExp::*relK*T77A. Cultures were grown overnight in LB media supplemented with Amp at 37°C with shaking at 180 rpm. Secondary inoculation was done with one percent of primary inoculum and further grown for 3-4 hrs (OD_600nm_ ∼ 0.6-0.8). Cultures were then diluted to an OD_600nm_ of 0.5, and 100 µL was aliquoted in triplicates onto the 96-well plate. IPTG was then added to induce protein expression. OD_600nm_ was recorded at an interval of 30 mins for 4 hrs to monitor growth with and without the inducer. Data was plotted as a function of growth over time. Viable counts were determined at 4 hr post induction by drop plating serial dilutions on LB agar plates containing antibiotic.

For co-expression studies, *E. coli* BL21 cells were co-transformed with the plasmids expressing RelJK_WT_ or RelJK_Mut_T77A and grown overnight in LB media supplemented with Amp, Kan (30 µg/mL) and Glc at 37°C with shaking at 180 rpm. Secondary inoculation was done with one percent of primary inoculum in medium without Glc and grown till OD_600nm_ ∼ 0.4-0.6. Cultures were then induced with ATc and Ara for 4 hrs. For CFU determination, cultures were serially diluted and drop plated on LB agar plates containing relevant supplements. Statistical analysis was done using one-way analysis of variance (ANOVA) using GraphPad Prism 9.0 software.

## Results

### *In silico* prediction of potential post-translational modifications of *M. tuberculosis* TA modules

Recently, we utilized a bioinformatic approach to report extensive phosphorylation in mycobacterial signaling proteins (39). Using the same strategy with newer tools, we investigated the potential of *M. tuberculosis* TA proteins to undergo post-translational modifications. We used MusiteDeep for PTM prediction, an online resource providing a deep-learning framework for protein post-translational modification site prediction and visualization (40–42). The predictor only uses protein sequences as input and no complex features are needed resulting in a realistic prediction for proteins (41). We analyzed 128 *M. tuberculosis* toxin and antitoxin proteins belonging to 64 paired TA modules of Vap, Maz, Rel, Par and Hig families, and grouped them in nine different PTM categories (Fig. 1A and 1B). Remarkably, the percentage of PTM by phosphorylation was significantly higher than other predicted PTMs (Fig. 1A and 1B) suggesting that phosphorylation of TA proteins may be a ubiquitous regulatory mechanism to modulate TA function. Interestingly, with the exception of 3 antitoxins (VapB3, VapB9 and MazE3), all antitoxins showed putative phosphosites. In contrast, 25 % of the toxins analyzed were not positive for phosphorylation. For smaller TA families such as Par and Hig, all TA proteins were amenable to Ser/Thr phosphorylation. These results clearly highlight extensive *O*-phosphorylation of *M. tuberculosis* toxin and antitoxin proteins.

**Figure 1.**
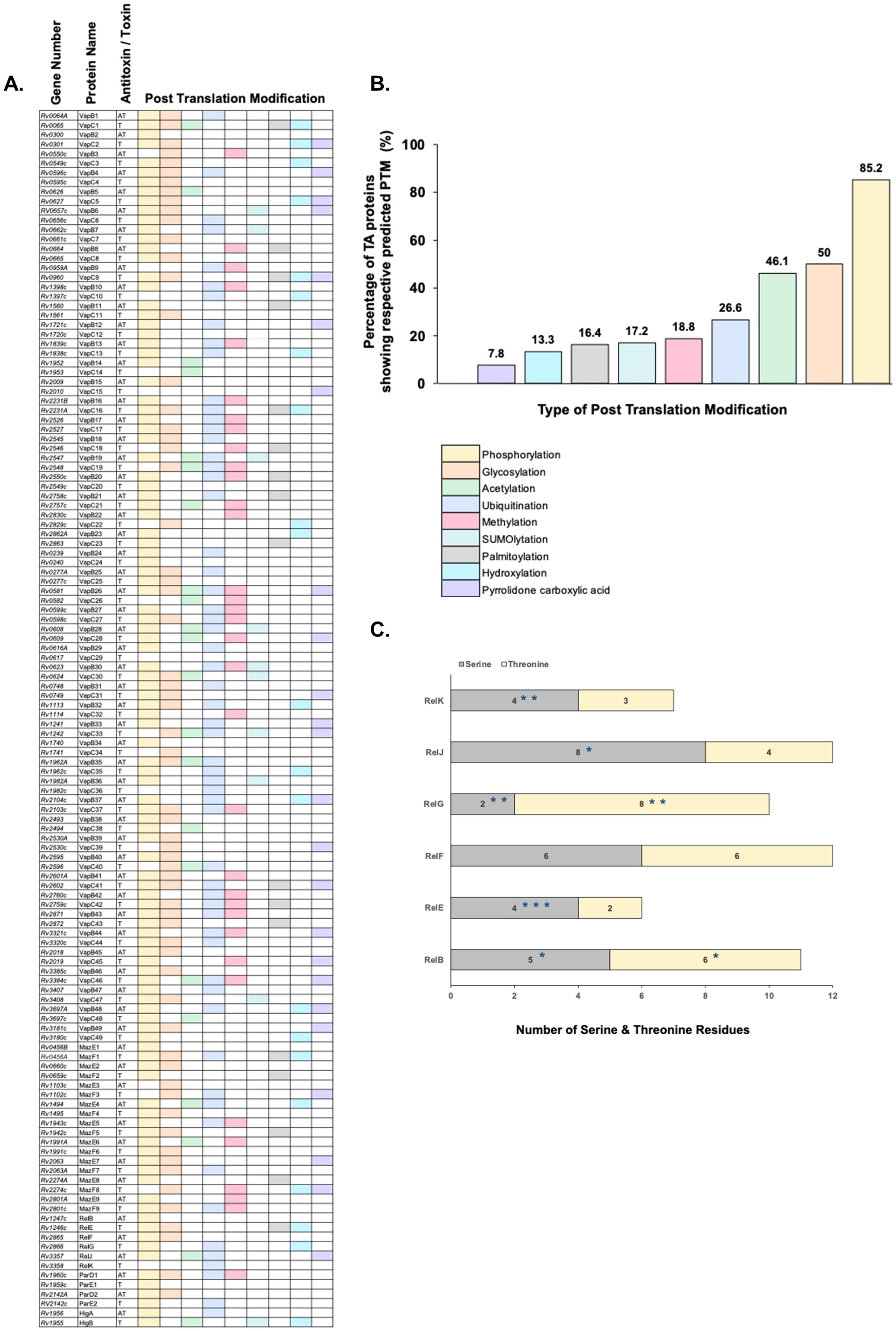
*In silico* analysis of predicted PTMs for *M. tuberculosis* TA proteins. **(A)** Distribution of nine different PTMs amongst the 128 *M. tuberculosis* Antitoxin (AT) and Toxin (T) genes as predicted by MusiteDeep software (https://www.musite.net/). The gene numbers and names for respective TA genes/proteins from *M. tuberculosis* H37Rv strain are listed. **(B)** The TA proteins showing a potential PTM were grouped together, and percentage is plotted to depict the % of TA proteins showing a respective PTM. **(C)** Potential Ser/Thr phosphosites in Rel TA family predicted by NetphosBac 1.0. Numbers in the stacked bar graph denote the total number of serine (grey) and threonine residues (yellow) in Rel TA proteins. Of the available Ser/Thr sites, the number of potential phosphosites predicted by NetphosBac 1.0 (https://services.healthtech.dtu.dk/services/NetPhosBac-1.0/) are marked with an Asterisk *.

Based on these results, we subjected the RelBE TA proteins to an additional screening for potential phosphosites using NetPhosBac 1.0 software. While MusiteDeep predicts PTMs, this tool is based on an artificial neural networking model that predicts bacteria specific *O*-phosphorylation (43). It should be noted that NetPhosBac considers the full-length of the protein irrespective of any specific domain regions, and only reports phosphosites with scores > 0.5. The total number of serine and threonine residues for each Rel protein are depicted in Figure 1C, and the predicted number of phosphosites are marked as an asterisk. In our analysis, we determined that all members of the Rel family except RelF are likely to be modified by *O-*phosphorylation. Overall, the propensity of potential serine phosphorylation was higher than threonine modification. It is noteworthy that Rel toxins seem more susceptible to STPK-mediated phosphorylation as compared to Rel antitoxins. Taken together, computational analyses revealed first, that *M. tuberculosis* TA modules are amenable to PTMs and second, that modification by *O-*phosphorylation may be a bona fide mode of TA regulation.

### PknK interacts with Rel toxin and antitoxin proteins *in vivo*

*M. tuberculosis* STPKs are established regulators of diverse cellular processes ranging from regulation of growth, cell division, membrane biogenesis, metabolism and pathogenesis (44). Considering that PknK plays a key role in stress-induced growth adaptation, we explored the role of PknK in phosphorylation of *M. tuberculosis* Rel TA proteins.

The M-PFC assay (37) was used to detect PknK interactions with either Rel toxin or antitoxin proteins in *M. smegmatis* cells. Previous results suggest that RelB and RelE directly bind, with interaction likely occurring between the C termini of both proteins (19). Additionally, Korch et al., observed that the toxicity of RelE was dependent upon a free C-termini, as RelE-C terminal fusion proteins were incapable of inhibiting mycobacterial growth, even in the absence of RelB. Directed by these results, we tested whether Rel antitoxin-_[F1,2]-N_ fusions or Rel toxin-_[F3]-N_ fusions interact with PknK-_[F1,2]-N_ or PknK-_[F3]-N_ fusion proteins in *M. smegmatis.* Controls included *M. smegmatis* co-transformed with the empty vectors pUAB100/pUAB200 (GCN4_[F1,2]_/GCN4_[F3]_) as positive control and the empty vector pUAB200/RelB_[F1,2]-N_ or with the empty vector pUAB300/RelE_[F3]-C_ as negative controls. As seen in Figures 2A and 2B, all antitoxin- or toxin-PknK co-transformations resulted in growth; however, the growth of RelE/G/K-PknK co-transformants was less robust, likely due to the toxin activity in absence of the antitoxin (Fig. 2B). Furthermore, when grown in the presence of TRIM and thus requiring protein interaction for survival, we only observed reproducible growth of *M. smegmatis* for the following protein pairs: RelJ_[F1,2]-N_/PknK_[F3]-N_, RelE_[F3]-N_/PknK_[F1,2]-N_, RelK_[F3]-N_/PknK_[F1,2]-N_ and the positive controls GCN4_[F1,2]_/GCN4_[F3]_ and RelB_[F1,2]-N_/RelE_[F3]-N_ (Fig. 2A and 2B). These results indicate that PknK selectively interacts with RelE and RelK toxins, and the RelJ antitoxin protein *in vivo*.

**Figure 2.**
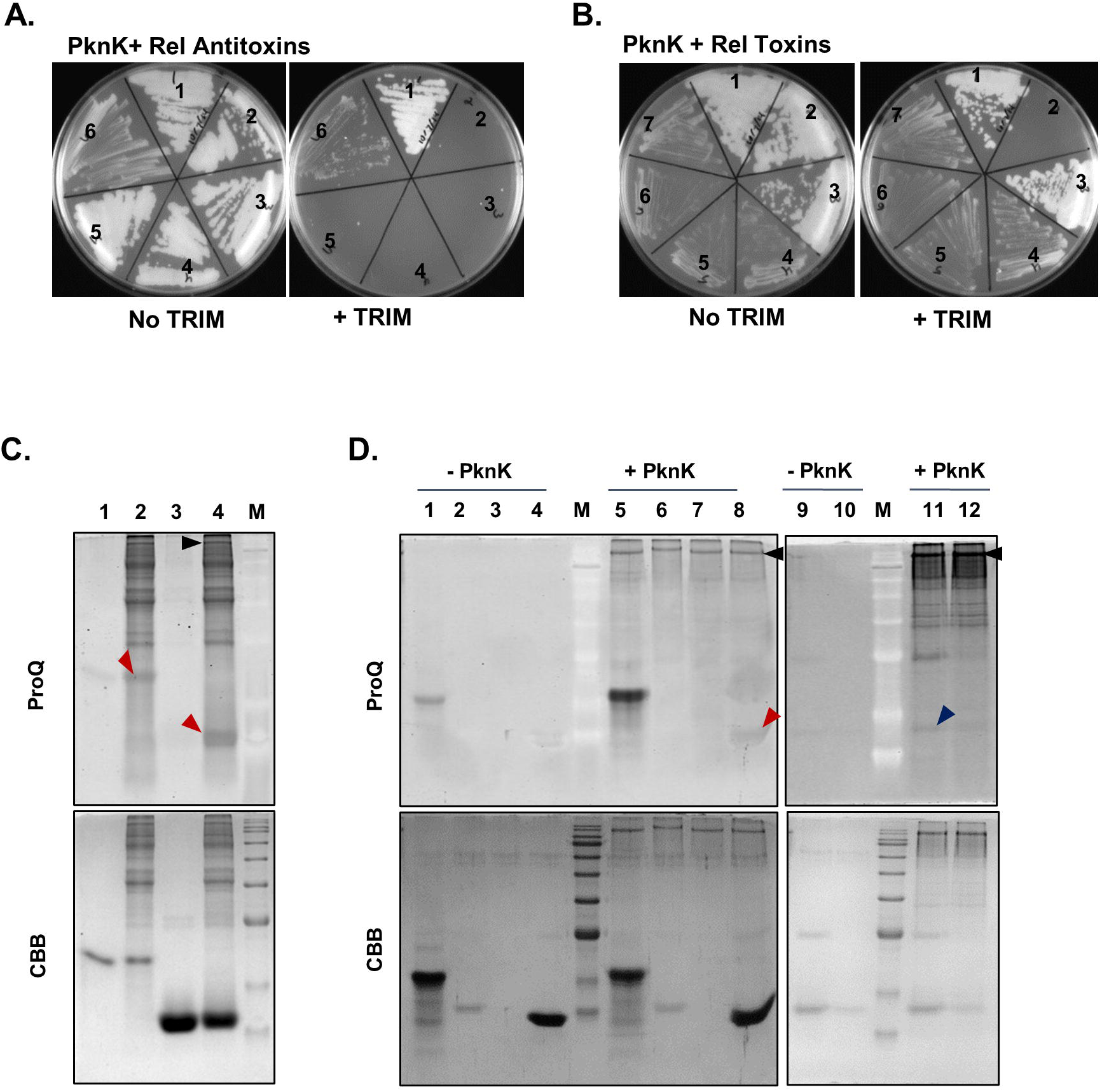
Rel TA proteins interact with PknK *in vivo* and *in vitro*. M-PFC assay depicting *in vivo* protein interactions between PknK and **(A)** Rel Antitoxins. *M. smegmatis* was co-transformed with positive control pUAB100 + pUAB200 (section 1), negative control RelB_[F1,2]-C_/pUAB200 (section 2), RelB_[F1,2]-N_/PknK_[F3]-N_ (section 3), RelB_[F1,2]-C_/PknK_[F3]-N_ (section 4), RelF_[F1,2]-N_/PknK_[F3]-N_ (section 5), RelJ_[F1,2]-N_/PknK_[F3]-N_ (section 6), and with **(B)** Rel toxins. *M. smegmatis* was co-transformed with positive control pUAB100 + pUAB200 (section 1), negative control RelE_[F3]-C_/pUAB300 (section 2), RelB_[F1,2]-N_/RelE_[F3]-N_ (section 3), RelE_[F3]-N_/PknK_[F1,2]-N_ (section 4), RelE_[F3]-C_/PknK_[F1,2]-N_ (section 5), RelG_[F3]-N_/PknK_[F1,2]-N_ (section 6), RelK_[F3]-N_/PknK_[F1,2]-N_ (section 7). Transformants were streaked on 7H11-Kan-Hyg plates and 7H11-Kan-Hyg-TRIM plates to select for protein-protein interactions. **(C)** *In vitro* kinase assay of wild type PknK with RelJ Antitoxin. All gels are stained with ProQ™ Diamond phosphostain on the top panel and CBB on the bottom panel. MBP was used as positive control. Lanes 1 and 2 show MBP without and with PknK, respectively. Lanes 3 and 4 show RelJ alone and with PknK, respectively. Red arrow on the top gel panel shows phosphorylation of MBP (Lane 2) and RelJ by PknK (Lane 4). **(D)** *In vitro* kinase assay of PknK with Rel Toxins (RelE, RelG, RelK). Lanes 1 and 5 show MBP without and with PknK, respectively. RelE, RelG and RelK alone (Lanes 2-4) and with PknK (Lanes 6-8), Red arrow on the top gel panel shows phosphorylation of RelK by PknK (Lane 8). Kinase assay of RelE and RelG with PknK (Lanes 11 and 12), Blue arrow on the top gel panel shows phosphorylation of RelE by PknK. Black arrows indicate phosphorylated PknK.

### Rel toxin and antitoxin proteins undergo *in vitro O*-phosphorylation by PknK

To investigate if protein-protein interactions between PknK and Rel TA proteins culminated into a PTM such as phosphorylation, we performed *in vitro* kinase assays with purified recombinant Rel proteins and wild type or phosphorylation defective PknK. To ensure the validity of all kinase assays, we confirmed the functional activity of purified PknK (autophosphorylation and transphosphorylation) using Myelin Basic Protein (MBP) as a general kinase substrate that also served as a positive control in our assays (Fig. 2C).

For kinase assays, positive reactions were visualized by ProQ™ Diamond Phosphoprotein Gel Stain that specifically stains phosphorylated S/T/Y amino acids (45). Rel proteins were tested for basal level of background phosphostaining in absence of the kinase. While no signal was observed with RelB and RelF antitoxins (data not shown), both MBP and RelJ showed change in signal intensities in the presence of wild type PknK as compared to proteins alone in ProQ-stained gels (Fig. 2C, Lanes 2 and 4 vs Lanes 1 and 3). Densitometric analysis using the ImageJ software revealed a ∼ 4-fold increase in the intensity of the phosphostain in reactions containing RelJ and PknK. With respect to toxin proteins, a ∼ 3.4-fold change in phosphosignal intensity was observed for RelK in presence of PknK as compared to RelK alone (Fig. 2D, compare Lanes 4 and 8). These observations suggest that both cognate TA proteins, RelJ and RelK are substrates for PknK-mediated phosphorylation. There was no phosphosignal observed for RelE and RelG proteins (Fig. 2D, Lanes 6 and 7). We noticed that the Rel toxins were susceptible to aggregation. Despite equal loading, the protein amounts observed in the CBB stained gel were considerably lower for RelE and RelG than RelK (Fig. 2D; Lanes 2 and 3, lower panel). Thus, we repeated assays with fresh preparations of RelE and RelG (Fig. 2D; *right*) and observed a faint signal for phosphoprotein staining with RelE in the presence of PknK and ATP (Fig. 2D, compare Lanes 9 and 11). There was no difference in signal intensities in case of RelG toxin.

To confirm that the phosphorylation signal observed for RelE, RelK and RelJ is PknK-specific, we performed the same assay in the presence of phosphorylation-defective PknK K55M mutant protein. As expected, there was no signal observed for the RelE, RelK or RelJ substrate proteins when incubated with the mutant PknK protein (data not shown). Based on these results, we concluded that toxins, RelE and RelK and antitoxin, RelJ are substrates for phosphorylation by PknK.

### Kinetics of PknK-mediated phosphorylation of Rel TA proteins

To determine the phosphorylation rates for RelE and RelJK proteins, we performed time kinetic analysis. The toxin and antitoxin proteins were incubated alone or with PknK for 0, 15, 45, 60 minutes and relative phosphorylation levels were calculated using the ImageJ software (Fig. 3). The trend of phosphorylation was similar for RelJ and RelK proteins attaining a maximum signal at 15 mins, followed by a consistent but insignificant decrease in the signal by 60 mins. RelE displayed a distinctively different pattern with a maximum phosphosignal observed at the 30- and 45 min after which there was a slight decrease in the signal intensity over 60 mins. We also determined the strength of the phosphorylation signal after ATP removal, over more extended time points upto 24 hrs and observed that the phosphosignal was stable.

**Figure 3.**
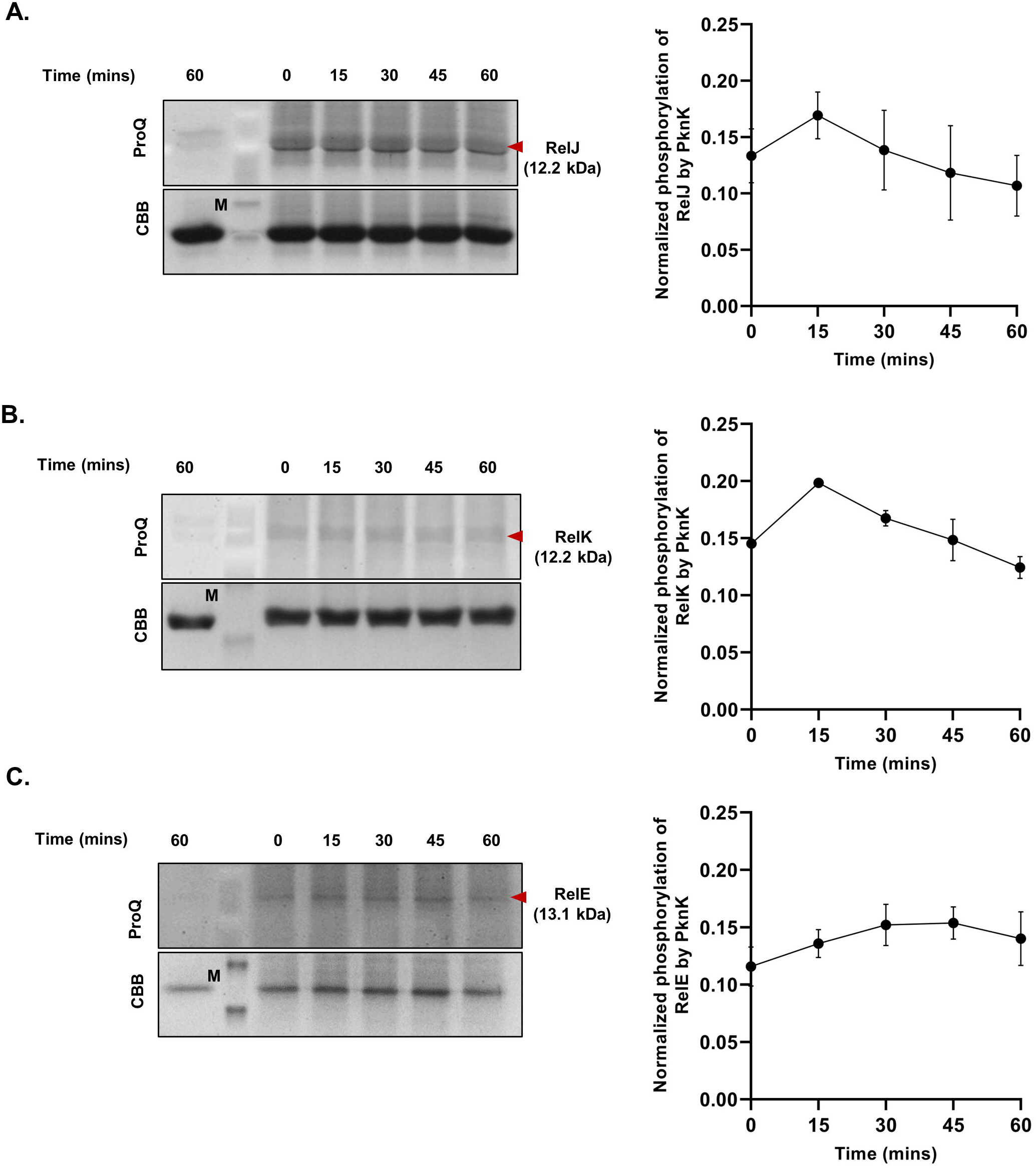
Time kinetics of PknK-mediated phosphorylation of Rel TA proteins. **(A)** RelJ antitoxin **(B)** RelK toxin and **(C)** RelE toxin. In all images, Lane 1 shows the substrates alone. Time course was done at the indicated time points with PknK and substrate i.e., Antitoxin or Toxin (indicated by red arrows). All gels are stained with ProQ™ Diamond on the top panel and CBB on the bottom panel. One representative gel picture is shown. Data is plotted as Mean ± SD from three independent experiments.

### Rel antitoxin proteins exhibit differential oligomerization

Crystal structure analysis of Rel TA proteins revealed that the antitoxin and toxin bind in a 1:1 ratio, with the heterodimer forming a tetramer through interactions occurring at the antitoxin-antitoxin interface (46). To gain insight into the structural organization of the TA proteins, we analyzed their individual electrophoretic profiles on an 8% native PAGE gel. As shown in Fig. 4A, the antitoxins have characteristically distinct oligomerization profiles. The oligomeric forms of the antitoxins appear to be inherently more stable as we were not able to observe the monomeric forms of the antitoxins. Strikingly, a ladder of oligomeric forms of RelB was observed that resulted in higher order structures having a different size as compared to RelF and RelJ tetramers suggesting that RelB forms extended structures that are unique. RelK toxin has a pI of 6.5, whereas RelE and RelG toxins have a highly basic pI (9.6-11.0) and thus, were not resolved on a native PAGE.

**Figure 4.**
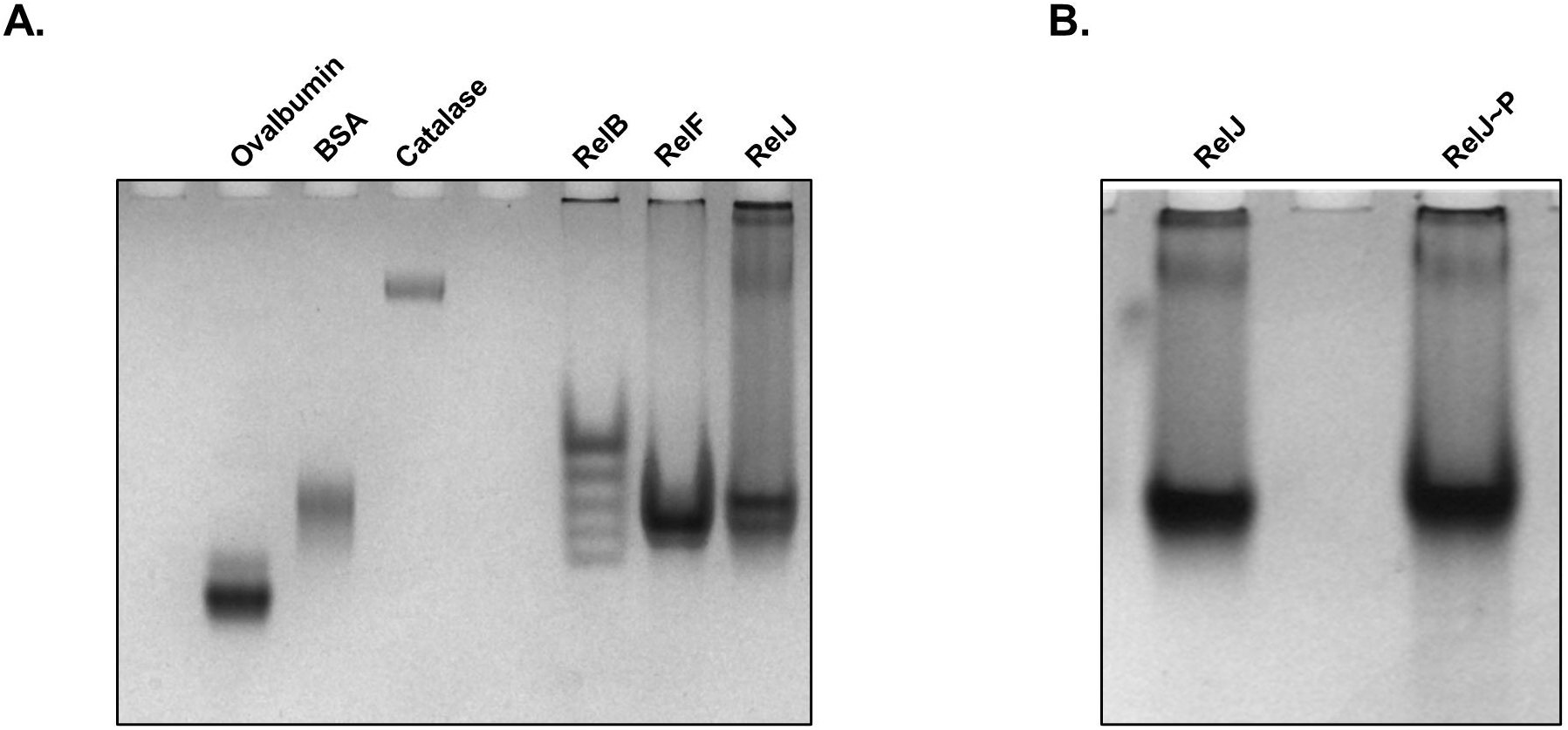
Oligomerization pattern of Rel TA proteins. Native gels were run to check the **(A)** oligomerization pattern of Antitoxin proteins (RelB, RelF, RelJ) and **(B)** to determine the effect of phosphorylation on oligomerization of RelJ antitoxin. Commercially purified native proteins such as Ovalbumin protein (MW ∼ 43 kDa), Bovine serum albumin protein (MW∼ 67 kDa), and Catalase protein (MW ∼240 kDa) were used as native PAGE molecular weight markers.

To determine whether phosphorylation or interaction of RelJ with PknK affected its oligomeric pattern, RelJ was incubated with PknK as done for the kinase assays and resolved on an 8% native PAGE. As evident from Fig. 4B, there is no observed change in the self-assembly of RelJ antitoxin with or without phosphorylation suggesting that the PTM is not in the oligomeric interface of the RelJ protein.

### Mass spectrometry reveals extensive phosphorylation of the RelJK TA module by PknK

To identify the specific amino acid phosphorylated by PknK, the phosphorylated TA proteins were subjected to LC-MS/MS analysis. As seen in Figure 5, MS spectra revealed a single phosphorylated site in RelK, identified as Threonine at position 77 (Fig. 5A). Additionally, several PknK-mediated phosphorylation sites were identified in RelJ antitoxin at positions 21 (Thr), 55 (Ser) and 74 (Ser) of the RelJ polypeptide chain (Fig. 5B).

**Figure 5.**
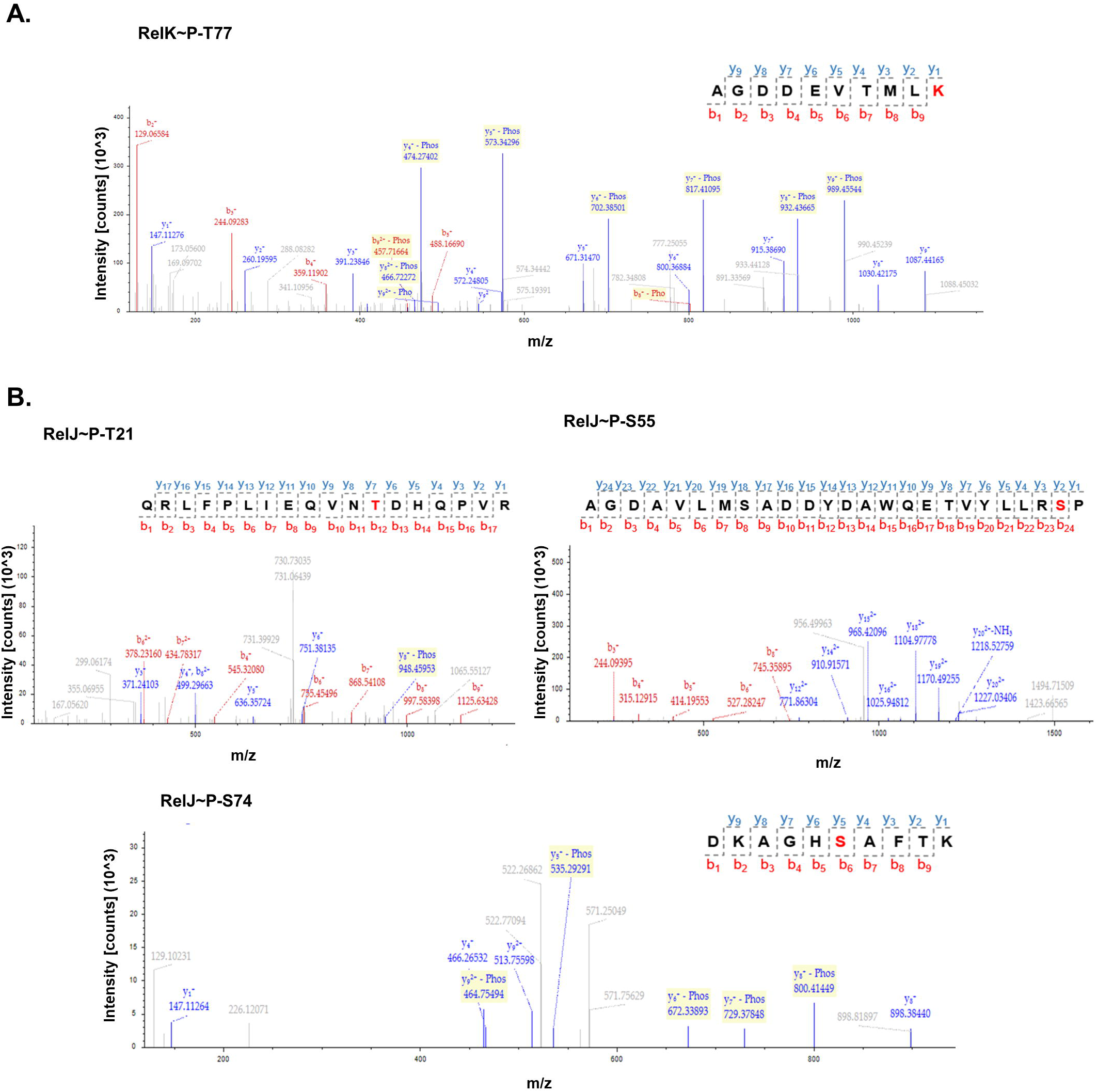
LC-MS/MS analysis for identification of phosphorylated residues. Mass spectra analysis for **(A)** RelK toxin and **(B)** RelJ antitoxin proteins. The respective peptides with the phosphorylated residue (highlighted in red) are shown in the inset. Peptide spectrum match and protein false discovery rate were set to 0.01 FDR to obtain high confidence hits.

### Phosphorylation of Thr77 does not affect the cytotoxic effect of RelK toxin

Of all the Rel toxins, the RelK toxin is known to exert a modest cytotoxic effect on growth in *E. coli* (18) and *M. smegmatis* (46). To understand the functional effects of RelK∼P, we replaced Thr77 in RelK with alanine by site directed mutagenesis as described in the methods. The resulting mutant protein RelK_Mut_T77A was confirmed to be phosphorylation deficient by LC-MS/MS analysis.

We compared the growth profiles of *E. coli* cells overexpressing RelK_WT_ and RelK_Mut_T77A in presence and absence of the inducer IPTG. Cell lysates were also analyzed to eliminate any significant differences in the levels of wild type and mutant toxins in our experimental set up (data not shown). Uninduced cells did not exhibit any defect in growth suggesting that there is no leaky expression of the toxins (Fig. 6A). In induced cells, both the RelK_WT_ and RelK_Mut_T77A toxins negatively affected growth post induction. In fact, the RelK_Mut_T77A displayed significantly reduced growth than the induced RelK_WT_ *E. coli* cells in liquid cultures (Fig. 6A). These results were replicated when the assay was done on solid medium. The RelK_Mut_T77A mutant toxin grew poorly on solid agar plates (Fig. 6B). The results suggest that neither Thr77 nor its phosphorylation are essential for the catalytic activity of the toxin, and that the mutant form of the toxin is more cytotoxic than the wild type protein.

**Figure 6.**
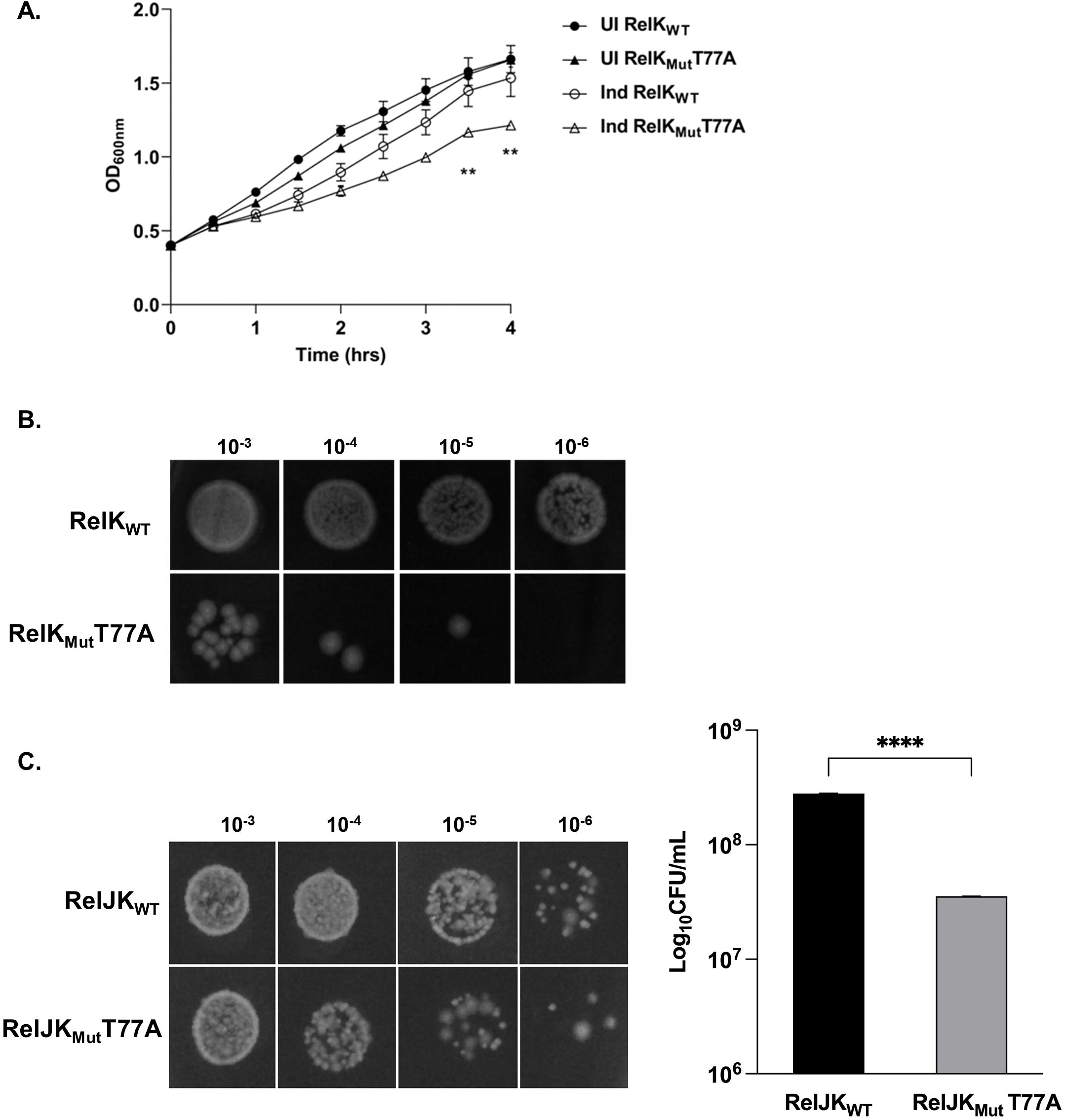
Cytotoxic effect and neutralization of RelK_WT_ and RelK_Mut_T77A toxins in *E. coli*. **(A)** Plot of OD_600_ versus time showing the cytotoxic effect of RelK_WT_ and RelK_Mut_T77A in *E. coli*. Open circle and triangle denote RelK_WT_ and RelK_Mut_T77A overexpressing cells, respectively that are induced (Ind) with 1.0 mM IPTG. Growth was compared with respective uninduced (UI) cells (filled circle and triangle). Data is plotted as Mean ± SD from three technical replicates. ** represents *p<0.001* for the difference in growth of cells overexpressing RelK_WT_ versus the RelK_Mut_T77A mutant protein under inducing conditions. **(B)** Viable counts of *E. coli* cells overexpressing RelK_WT_ or RelK_Mut_T77A induced with IPTG for 4 hrs, followed by plating of serial dilutions on the LB-Amp plates. **(C)** *E. coli* cells co-expressing wild type RelJ antitoxin with RelK_WT_ or RelK_Mut_T77A toxins were induced with ATc and Ara for antitoxin and toxin production, respectively. Serial dilutions were drop plated on solid medium containing Kan, Amp and Glc to obtain viable counts. Bar graph showing viable counts obtained in cells co-expressing RelJK_WT_ versus RelJK_Mut_T77A is presented. The decrease in viability of RelJK_Mut_T77A co-expressing cells indicates inefficient neutralization of the mutant toxin by the RelJ antitoxin. **** represents *p<0.00001* for the difference in growth of cells co-expressing RelK_WT_ or RelK_Mut_T77A toxins with wild type RelJ antitoxin.

### Rescue studies of RelJK_WT_ and RelJK_Mut_T77A in *E. coli*

Neutralization of the RelK toxin by its cognate antitoxin RelJ has been described earlier in *E. coli* (18) and *M. smegmatis* (19). The amount of free toxin available in the cell is dictated by its cognate antitoxin which binds to and neutralizes toxin activity. Thus, we wanted to determine if insufficient neutralization of the mutant toxin compared to the wild type toxin by RelJ antitoxin could be a contributory factor in the growth-related phenotype observed in Fig. 6A. Towards this, we co-expressed RelJ and RelK_WT_ or RelK_Mut_T77A in *E. coli* as described previously (18). The co-transformants were induced with Arabinose (Ara) and Anhydrotetracycline (ATc) to induce the expression of both toxin and antitoxin genes, respectively. As shown in Fig. 6C, the RelJK_Mut_T77A co-expressing cells showed decreased growth as compared to cells co-expressing RelJK_WT_. Assuming that complete toxin neutralization resulted in maximal growth of the RelJK_WT_ co-transformed cells (set at 100 %), there was a 13 % decrease in growth of RelJK_Mut_T77A co-transformed cells (Fig. 6C) indicating partial neutralization of the mutant toxin. Although Thr77 is not directly required for activity of the RelK toxin as seen in Figure 6A, its absence impacts toxin neutralization by RelJ antitoxin. While these findings clearly implicate Thr77 at the antitoxin binding region, the role of its phosphorylation in antitoxin binding remained unclear.

### Phosphorylation of the RelK toxin interferes with its binding to RelJ antitoxin

Guided by our results, we sought to investigate how phosphorylation of RelK affects its interaction with cognate antitoxin RelJ. Towards this objective, we incubated RelJK_WT_ and RelJK_Mut_T77A for 1 hr and resolved the complexes on an 8 % native gel. As seen previously, the toxin proteins alone did not resolve on the gel; however, incubation of wild type proteins, RelK_WT_ with RelJ in equimolar ratio produced a significant shift in the mobility of RelJ (Fig.7A, Lane 3). Of significance, the majority of RelJ was observed in a bound state, as demonstrated by the shift in molecular size. By contrast, only a slight shift in RelJ mobility was observed when incubated with increasing concentration of RelK_Mut_T77A protein indicating poor binding of the mutant RelK protein with the RelJ antitoxin (Fig.7A, Lanes 4 and 5).

**Figure 7.**
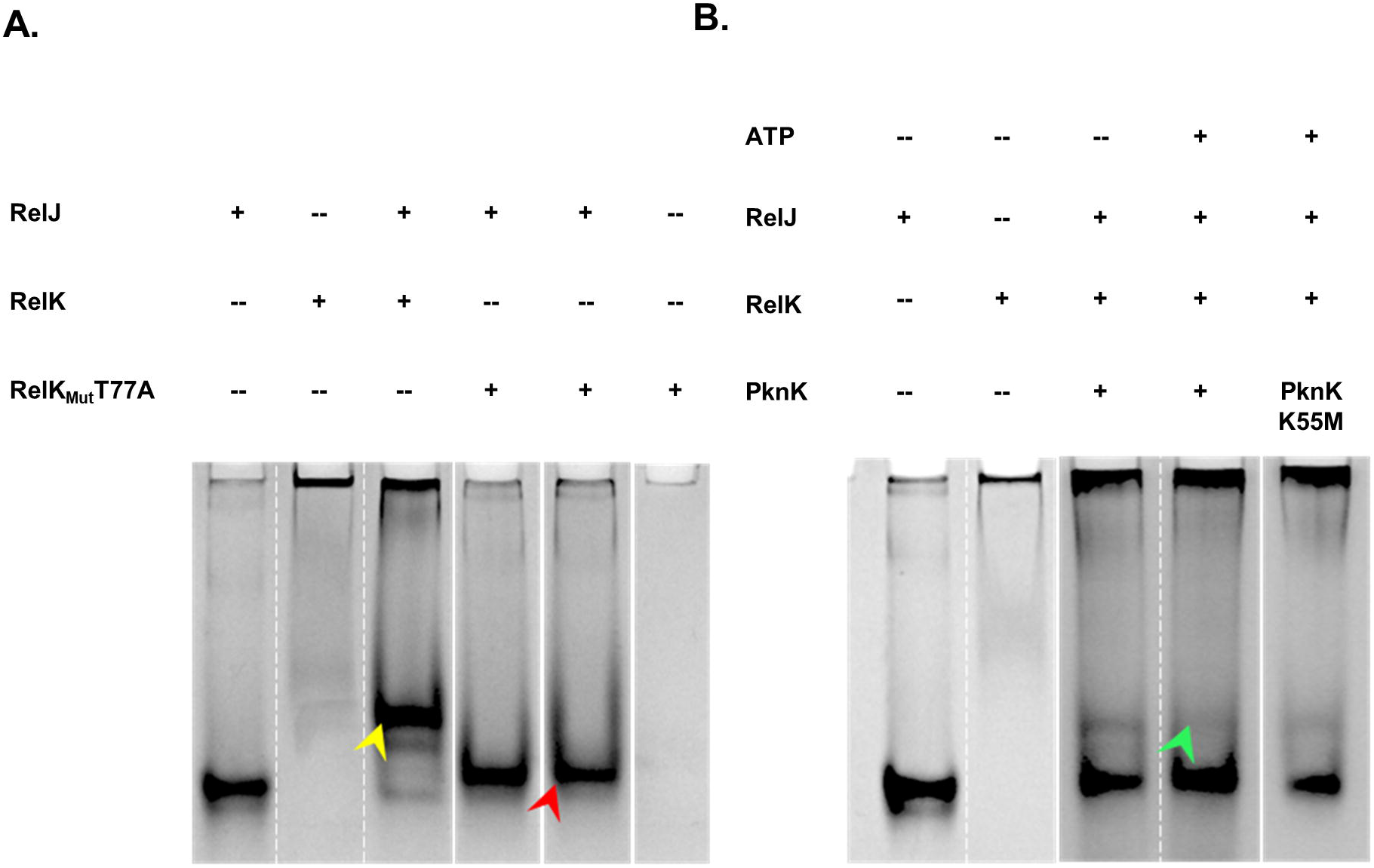
Effect of phosphorylation on interaction of wild type and mutant RelJK proteins on 8 % Native PAGE. **(A)** Binding of RelJ with RelK_WT_ or RelK_Mut_T77A proteins. Lanes 1, 2 and 6 are RelJ, and RelK_WT_ and RelK_Mut_T77A alone, respectively. RelJ was incubated with RelK_WT_ in equimolar ratio (Lane 3) and with RelK_Mut_T77A in increasing molar ratios (Lane 4 and 5). **(B)** Effect of phosphorylation on RelJK interaction. Lanes 1 and 2 are RelJ and RelK alone, respectively. RelK was incubated with RelJ in equimolar ratio with PknK without ATP (Lane 3) and with ATP (Lane 4), and with PknK K55M (Lane 5). Yellow arrow indicates the shift observed upon binding of RelJ with RelK wild type proteins. On the contrary, only a slight shift was observed when RelJ was incubated with RelK_Mut_T77A (red arrow) protein suggesting that Thr77 residue plays a role in the binding interface and that its phosphorylation (green arrow) interferes with binding of RelK to RelJ antitoxin.

We further compared RelJK complex formation in unphosphorylated and phosphorylated states. Expectedly, in the presence of PknK and ATP, the complex formation was significantly reduced as indicated by the green arrow (Fig. 7B, Lane 4). We observed comparable shifts in RelJ mobility when RelJK proteins were incubated with PknK minus ATP or with the phosphorylation defective PknK mutant (Fig. 7B, Lanes 3 and 5). The absence of any shift in Lane 4 of Fig. 7B confirmed that PknK-mediated phosphorylation interferes with RelJK complex formation. Cumulatively, the results presented in Figures 6 and 7 elucidate an integral role of Thr77 residue of RelK toxin in facilitating its binding to the RelJ antitoxin. More importantly, these findings establish toxin phosphorylation as means to interfere with toxin-antitoxin interaction resulting in inefficient toxin neutralization and increased cytotoxicity.

## Discussion

Emerging evidence has implicated a role for the toxin-antitoxin modules in the initiation, development, and establishment of the persistence phenotype in *Mycobacterium tuberculosis.* Although functional characterization of multiple Type II TA systems has been reported, there is little evidence to describe the underlying mechanisms that regulate the coordinated expression of mycobacterial TA systems. In this study, we have investigated the role of post-translational modifications (PTMs) in the regulation of *M. tuberculosis* Type II TA modules. Using biochemical and computational biology approach, we establish that *O*-phosphorylation of TA proteins by *M. tuberculosis* Ser/Thr protein kinases is a distinct and novel mode of regulating toxin-antitoxin interaction.

Protein post-translational modifications are ubiquitous events that have important functional consequences such as altering enzyme activity, inhibiting DNA binding activity or altering protein conformation (47). PTMs promise to offer exquisite control of the enzymatic activities, which in the case of TA modules and their established roles in mycobacterial persistence, is crucial to avoid spurious initiation of persister formation. So far, very little is known about PTM of TA modules, thus understanding the role of PTMs in regulation of TA function is of significant interest. *In silico* analyses revealed that TA modules are amenable to distinctly different post-translational modifications with phosphorylation leading the list followed by glycosylation. Recently, Yu et al., reported a mechanism of toxin neutralization via phosphorylation (29). TakA-TglT encoded by *Rv1044-Rv1045* constitute a unique TA module wherein the antitoxin TakA is an atypical kinase that phosphorylates the TglT toxin resulting in its neutralization (29). It is noteworthy that *M. tuberculosis* is equipped with distinct eukaryotic-like protein kinases (32, 33) and that TakA has very little similarity with these proteins, thus providing the rationale to investigate the role of STPKs in the regulation of TA modules. During the preparation of this manuscript, Frando et al., reported widespread *O*-phosphorylation of the *M. tuberculosis* H37Rv proteome (48). While their data implicated multiple STPKs in TA regulation, this study is the first instance that provides direct biochemical evidence of post-translational modification of mycobacterial Type II Rel TA modules via PknK highlighting a novel underlying regulatory mechanism to modulate TA function.

The RelBE toxin-antitoxin system has been extensively studied in *E. coli* wherein the RelE toxin is activated under the stringent response and specifically cleaves ribosome associated mRNAs, thereby reducing the global level of translation (49). Likewise, the *M. tuberculosis* RelBE homologs are functional mediators of persistence involved in stress-induced translational control (34) with the toxins exhibiting differential cytotoxic effects (46). While the *M. tuberculosis* RelE toxin is an acutely toxic, mRNA interferase, that in the absence of RelB antitoxin, induces a viable but dormant phenotype to a large proportion of the population (19), the RelK toxin is moderately cytotoxic (46). These inherent differences stem from the structural and sequence divergence of these proteins such that RelBE (RelBE1) and RelFG (RelBE2) are grouped together in the RelBE sub family within the Rel superfamily while RelJK (RelBE3) is considered a part of the *E. coli* YefM/YoeB system (46). Resolution of the RelK crystal structure affirmed that the catalytic mechanism of the RelK toxin is closer to the YoeB toxin than the RelE or RelG toxins (46). At present, there is no information regarding the cellular targets or substrates of the RelK toxin. However, a YoeB toxin from *Agrobacterium tumefaciens* was recently shown to cleave both RNA and DNA with similar efficiency (50) suggesting that YoeB-like toxins can act as non-specific, ribosome independent nucleases. Whether this is true for *M. tuberculosis* RelK toxin remains to be investigated. In accordance with their role in mycobacterial persistence, transcripts specific for *relE* and *relK* toxins, and *relF* antitoxin have been detected in *M. tuberculosis*-infected human macrophages (19). Therefore, it is not surprising that these TA particularly *relFG* and *relJK* are significantly upregulated in response to antibiotic exposure (51), nitrogen starvation (34), oxidative stress (33) and in lungs of infected mice (51) and thus may be subjected to multiple levels of regulatory controls.

A key finding of this study is that PknK, a cytosolic STPK interacts with RelE and RelJK TA proteins and directs Ser/Thr phosphorylation at multiple sites. We detected three phosphosites on the RelJ antitoxin (Thr21, Ser55, Ser74) and a single phosphosite (Thr77) on the RelK toxin. Despite several attempts, phosphosites could not be determined for RelE toxin, probably due to consistent low yield and aggregation observed during toxin purification. It is apparent from the growth studies analyzing RelK_Mut_T77A that Thr77 is not required for toxin activity. However, the consistently observed higher toxicity obtained with cells overexpressing RelK_Mut_T77A compared to the wild type was surprising. We rationalized that the hyperactivity of the mutant toxin could stem from two possibilities, one that is directly linked to its catalytic activity and the other to its availability as a free toxin. Notably, cross neutralization of the Rel toxins in *M. smegmatis* has been reported (52). The authors demonstrated that RelB antitoxin could neutralize non-cognate RelG toxin in mycobacteria. (52). Thus, it is plausible that neutralization of the RelK_WT_ toxin by non-cognate, heterologous antitoxins of *E. coli* is responsible for the comparatively better survival of the cells overexpressing RelK_WT_ and that the same phenomenon is ineffective for the RelK mutant. This hypothesis was tested in co-expression studies where the cognate RelJ antitoxin enabled only partial rescue of cells overexpressing the RelK_Mut_T77A toxin. Based on these observations and *in vitro* binding studies, we propose that Thr77 in RelK is important for its interaction with the RelJ antitoxin. Furthermore, Thr77 is non-essential for toxin activity but plays a regulatory role via STPK-mediated post-translational modification.

The RelJK complex exists as a tetramer of two molecules of RelJ and RelK each (46), however the role of oligomerization in its function (if any) is unknown. We show that RelJ forms homo-oligomers *in vitro*, with the tetrameric form being the most prevalent. More importantly, as shown in Figure 4, PknK-mediated phosphorylation of the RelJ antitoxin did not alter its oligomeric pattern (Fig. 4B); however, it does certainly contribute towards inhibiting RelJK complex formation (Fig. 7). The role of the antitoxin is two-fold, first to bind and sequester the toxin and second, to autoregulate transcription of the TA operon. It is known that under stressful environments, the antitoxin is cleaved by proteases to free the toxin (13). Characterization of RelJ phosphorylation and its role in antitoxin stability/turnover or its autoregulatory function necessitates systematic mutational analysis and experimentation that are the focus of future research.

With clear benefits to prokaryotes, and correspondingly detrimental consequences to their host, a rapid, global TA-response could assist in antibiotic evasion and persistence maintenance. Our study sheds light on previously uncharacterized mechanisms related to mycobacterium persistence and has the potential to advance understanding of the current medical enigma of bacterial persistence facilitating development of new anti-tubercular therapies. We establish Ser/Thr phosphorylation of the toxin component as means to disrupt TA complex formation, rendering the toxin to inhibit growth and initiate persistence. The upregulation of *rel* and *pknK* transcripts under conditions of nitrogen starvation (34, 35) suggests that these cellular environments may be where PknK-mediated phosphorylation of Rel proteins is most physiologically relevant. We predict that other STPKs may similarly act to modify the TA proteins under different activating environments. If this is true, and the cross interactions between TA systems are extended to other mycobacterial TA modules, then it would result in a highly complex stress-response network, one that will undoubtedly influence *M. tuberculosis* resilience and pathogenic fitness.

## Acknowledgements.

We acknowledge CIF, UDSC for infrastructural support and Valerian Chem Pvt. Ltd., New Delhi for LC-MS/MS analysis. We are grateful to Prof. Adrie Steyn, USA for the M-PFC vectors. We also thank Prof. Deepak Saini, IISc, Bangalore and Dr. Lakshay Malhotra, SVC, New Delhi for their valuable inputs.

## Funding sources

This work was supported by the Department of Biotechnology, Govt. of India, DBT Grant No.BT/PR31937/MED/29/1404/2019 to VM. AG is thankful to DBT for JRF funding.

## Author contributions

SRS and AG planned and performed the experiments, analyzed the data, and wrote the manuscript; SK performed *in vivo* protein-protein interaction studies, AG provided project supervision support, VM conceived the study, provided project administration and supervision, analyzed data, and wrote the original manuscript.

## Notes

### Competing Interest Statement

The authors have declared no competing interest.

## References.

1. Bagcchi S. 2023. WHO’s Global Tuberculosis Report 2022. Lancet Microbe 4:e20.

2. Gomez JE, McKinney JD. 2004. M. tuberculosis persistence, latency, and drug tolerance. Tuberculosis 84:29–44.

3. Chai Q, Zhang Y, Liu CH. 2018. Mycobacterium tuberculosis: An Adaptable Pathogen Associated With Multiple Human Diseases. Front Cell Infect Microbiol 8:158.

4. Bigger J. 1944. Treatment of Staphyloeoeeal Infections with Penicillin by Intermittent Sterilisation. Lancet 497–500.

5. Kussell E, Kishony R, Balaban NQ, Leibler S. 2005. Bacterial Persistence. Genetics 169:1807–1814.

6. Wiuff C, Zappala RM, Regoes RR, Garner KN, Baquero F, Levin BR. 2005. Phenotypic tolerance: antibiotic enrichment of noninherited resistance in bacterial populations. Antimicrob Agents Chemother 49:1483–1494.

7. Moyed HS, Bertrand KP. 1983. hipA, a newly recognized gene of Escherichia coli K-12 that affects frequency of persistence after inhibition of murein synthesis. J Bacteriol 155:768–775.

8. Korch SB, Henderson TA, Hill TM. 2003. Characterization of the *hipA7* allele of *Escherichia coli* and evidence that high persistence is governed by (p)ppGpp synthesis. Mol Microbiol 50:1199–1213.

9. Keren I, Shah D, Spoering A, Kaldalu N, Lewis K. 2004. Specialized Persister Cells and the Mechanism of Multidrug Tolerance in *Escherichia coli*. J Bacteriol 186:8172–8180.

10. Germain E, Castro-Roa D, Zenkin N, Gerdes K. 2013. Molecular Mechanism of Bacterial Persistence by HipA. Mol Cell 52:248–254.

11. Amato SM, Orman MA, Brynildsen MP. 2013. Metabolic Control of Persister Formation in Escherichia coli. Mol Cell 50:475–487.

12. Hayes F. 2003. Toxins-Antitoxins: Plasmid Maintenance, Programmed Cell Death, and Cell Cycle Arrest. Science 301:1496–1499.

13. Gerdes K, Christensen SK, Løbner-Olesen A. 2005. Prokaryotic toxin–antitoxin stress response loci. Nat Rev Microbiol 3:371–382.

14. Hayes F, Van Melderen L. 2011. Toxins-antitoxins: diversity, evolution and function. Crit Rev Biochem Mol Biol 46:386–408.

15. Harms A, Brodersen DE, Mitarai N, Gerdes K. 2018. Toxins, Targets, and Triggers: An Overview of Toxin-Antitoxin Biology. Mol Cell 70:768–784.

16. Song S, Wood TK. 2020. Toxin/Antitoxin System Paradigms: Toxins Bound to Antitoxins Are Not Likely Activated by Preferential Antitoxin Degradation. Adv Biosyst 4:1900290.

17. Zhang S-P, Wang Q, Quan S-W, Yu X-Q, Wang Y, Guo D-D, Peng L, Feng H-Y, He Y-X. 2020. Type II toxin–antitoxin system in bacteria: activation, function, and mode of action. Biophys Rep 6:68–79.

18. Gupta A. 2009. Killing activity and rescue function of genome-wide toxin–antitoxin loci of *Mycobacterium tuberculosis*. FEMS Microbiol Lett 290:45–53.

19. Korch SB, Contreras H, Clark-Curtiss JE. 2009. Three *Mycobacterium tuberculosis* Rel Toxin-Antitoxin Modules Inhibit Mycobacterial Growth and Are Expressed in Infected Human Macrophages. J Bacteriol 191:1618–1630.

20. Frampton R, Aggio RBM, Villas-Bôas SG, Arcus VL, Cook GM. 2012. Toxin-Antitoxin Systems of Mycobacterium smegmatis Are Essential for Cell Survival. J Biol Chem 287:5340–5356.

21. Tripathi A, Dewan PC, Siddique SA, Varadarajan R. 2014. MazF-induced Growth Inhibition and Persister Generation in Escherichia coli. J Biol Chem 289:4191–4205.

22. Ramage HR, Connolly LE, Cox JS. 2009. Comprehensive Functional Analysis of Mycobacterium tuberculosis Toxin-Antitoxin Systems: Implications for Pathogenesis, Stress Responses, and Evolution. PLoS Genet 5:e1000767.

23. Sala A, Bordes P, Genevaux P. 2014. Multiple Toxin-Antitoxin Systems in Mycobacterium tuberculosis. Toxins 6:1002–1020.

24. Gupta A, Venkataraman B, Vasudevan M, Gopinath Bankar K. 2017. Co-expression network analysis of toxin-antitoxin loci in Mycobacterium tuberculosis reveals key modulators of cellular stress. Sci Rep 7:5868.

25. Sharma A, Sagar K, Chauhan NK, Venkataraman B, Gupta N, Gosain TP, Bhalla N, Singh R, Gupta A. 2021. HigB1 Toxin in Mycobacterium tuberculosis Is Upregulated During Stress and Required to Establish Infection in Guinea Pigs. Front Microbiol 12:748890.

26. Masachis S, Darfeuille F. 2018. Type I Toxin-Antitoxin Systems: Regulating Toxin Expression via Shine-Dalgarno Sequence Sequestration and Small RNA Binding, p. 171–190. *In* Storz, G, Papenfort, K (eds.), Regulating with RNA in Bacteria and Archaea. ASM Press, Washington, DC, USA.

27. Slayden RA, Dawson CC, Cummings JE. 2018. Toxin–antitoxin systems and regulatory mechanisms in Mycobacterium tuberculosis. Pathog Dis 76.

28. Yao J, Zhen X, Tang K, Liu T, Xu X, Chen Z, Guo Y, Liu X, Wood TK, Ouyang S, Wang X. 2020. Novel polyadenylylation-dependent neutralization mechanism of the HEPN/MNT toxin/antitoxin system. Nucleic Acids Res 48:11054–11067.

29. Yu X, Gao X, Zhu K, Yin H, Mao X, Wojdyla JA, Qin B, Huang H, Wang M, Sun Y-C, Cui S. 2020. Characterization of a toxin-antitoxin system in Mycobacterium tuberculosis suggests neutralization by phosphorylation as the antitoxicity mechanism. Commun Biol 3:216.

30. Dawson CC, Cummings JE, Starkey JM, Slayden RA. 2022. Discovery of a novel type IIB RELBE toxin-antitoxin system in *Mycobacterium tuberculosis* defined by co-regulation with an antisense RNA. Mol Microbiol 117:1419–1433.

31. Walsh C. 2006. Posttranslational modification of proteins: expanding nature’s inventory. Roberts and Company Publishers.

32. Cole ST, Brosch R, Parkhill J, Garnier T, Churcher C, Harris D, Gordon SV, Eiglmeier K, Gas S, Barry CE, Tekaia F, Badcock K, Basham D, Brown D, Chillingworth T, Connor R, Davies R, Devlin K, Feltwell T, Gentles S, Hamlin N, Holroyd S, Hornsby T, Jagels K, Krogh A, McLean J, Moule S, Murphy L, Oliver K, Osborne J, Quail MA, Rajandream M-A, Rogers J, Rutter S, Seeger K, Skelton J, Squares R, Squares S, Sulston JE, Taylor K, Whitehead S, Barrell BG. 1998. Deciphering the biology of Mycobacterium tuberculosis from the complete genome sequence. Nature 396:190–190.

33. Av-Gay Y, Everett M. 2000. The eukaryotic-like Ser/Thr protein kinases of Mycobacterium tuberculosis. Trends Microbiol 8:238–244.

34. Korch SB, Malhotra V, Contreras H, Clark-Curtiss JE. 2015. The Mycobacterium tuberculosis relBE toxin:antitoxin genes are stress-responsive modules that regulate growth through translation inhibition. J Microbiol 53:783–795.

35. Malhotra V, Okon BP, Satsangi AT, Das S, Waturuocha UW, Vashist A, Clark-Curtiss JE, Saini DK. 2022. Mycobacterium tuberculosis PknK Substrate Profiling Reveals Essential Transcription Terminator Protein Rho and Two-Component Response Regulators PrrA and MtrA as Novel Targets for Phosphorylation. Microbiol Spectr 10:e01354–21.

36. Malhotra V, Okon BP, Clark-Curtiss JE. 2012. Mycobacterium tuberculosis Protein Kinase K Enables Growth Adaptation through Translation Control. J Bacteriol 194:4184– 4196.

37. Singh A, Mai D, Kumar A, Steyn AJC. 2006. Dissecting virulence pathways of *Mycobacterium tuberculosis* through protein–protein association. Proc Natl Acad Sci 103:11346–11351.

38. Pelletier JN, Campbell-Valois F-X, Michnick SW. 1998. Oligomerization domain-directed reassembly of active dihydrofolate reductase from rationally designed fragments. Proc Natl Acad Sci 95:12141–12146.

39. Garg A, Khurana N, Chugh A, Verma K, Malhotra V. 2022. In silico evidence for extensive Ser/Thr phosphorylation of Mycobacterium tuberculosis two-component signalling systems. Curr Sci 123:1164.

40. Wang D, Zeng S, Xu C, Qiu W, Liang Y, Joshi T, Xu D. 2017. MusiteDeep: a deep-learning framework for general and kinase-specific phosphorylation site prediction. Bioinformatics 33:3909–3916.

41. Wang D, Liu D, Yuchi J, He F, Jiang Y, Cai S, Li J, Xu D. 2020. MusiteDeep: a deep-learning based webserver for protein post-translational modification site prediction and visualization. Nucleic Acids Res 48:W140–W146.

42. Wang D, Liang Y, Xu D. 2019. Capsule network for protein post-translational modification site prediction. Bioinformatics 35:2386–2394.

43. Miller ML, Soufi B, Jers C, Blom N, Macek B, Mijakovic I. 2009. NetPhosBac – A predictor for Ser/Thr phosphorylation sites in bacterial proteins. PROTEOMICS 9:116– 125.

44. Pereira SF, Goss L, Dworkin J. 2011. Eukaryote-like serine/threonine kinases and phosphatases in bacteria. Microbiol Mol Biol Rev 75:192–212.

45. Martin K, Steinberg TH, Cooley LA, Gee KR, Beechem JM, Patton WF. 2003. Quantitative analysis of protein phosphorylation status and protein kinase activity on microarrays using a novel fluorescent phosphorylation sensor dye. PROTEOMICS 3:1244–1255.

46. Miallau L, Jain P, Arbing MA, Cascio D, Phan T, Ahn CJ, Chan S, Chernishof I, Maxson M, Chiang J, Jacobs WR, Eisenberg DS. 2013. Comparative Proteomics Identifies the Cell-Associated Lethality of M. tuberculosis RelBE-like Toxin-Antitoxin Complexes. Structure 21:627–637.

47. Macek B, Forchhammer K, Hardouin J, Weber-Ban E, Grangeasse C, Mijakovic I. 2019. Protein post-translational modifications in bacteria. Nat Rev Microbiol 17:651–664.

48. Frando A, Boradia V, Gritsenko M, Beltejar C, Day L, Sherman DR, Ma S, Jacobs JM, Grundner C. 2023. The Mycobacterium tuberculosis protein O-phosphorylation landscape. Nat Microbiol 8:548–561.

49. Christensen SK, Gerdes K. 2003. RelE toxins from Bacteria and Archaea cleave mRNAs on translating ribosomes, which are rescued by tmRNA. Mol Microbiol 48:1389–1400.

50. McGillick J, Ames JR, Murphy T, Bourne CR. 2021. A YoeB toxin cleaves both RNA and DNA. Sci Rep 11:3592.

51. Singh R, Barry CE, Boshoff HIM. 2010. The Three RelE Homologs of *Mycobacterium tuberculosis* Have Individual, Drug-Specific Effects on Bacterial Antibiotic Tolerance. J Bacteriol 192:1279–1291.

52. Yang M, Gao C, Wang Y, Zhang H, He Z-G. 2010. Characterization of the Interaction and Cross-Regulation of Three Mycobacterium tuberculosis RelBE Modules. PLoS ONE 5:e10672.

